# PHENOTYPIC CHARACTERIZATION OF TWO NOVEL CELL LINE MODELS OF CASTRATION RESISTANT PROSTATE CANCER

**DOI:** 10.1101/2021.07.04.450352

**Authors:** Michael C. Haffner, Akshay Bhamidipati, Harrison K. Tsai, David M. Esopi, Ajay M. Vaghasia, Jin-Yih Low, Radhika A. Patel, Gunes Guner, Minh-Tam Pham, Nicole Castagna, Jessica Hicks, Nicolas Wyhs, Ruedi Aebersold, Angelo M. De Marzo, William G. Nelson, Tiannan Guo, Srinivasan Yegnasubramanian

**Affiliations:** Divisions of Human Biology and Clinical Research, Fred Hutchinson Cancer Research Center, Seattle, WA, USA; Department of Pathology, University of Washington, Seattle, WA, USA; Department of Pathology, Johns Hopkins University School of Medicine, Baltimore, MD, USA; Sidney Kimmel Comprehensive Cancer Center, Johns Hopkins University School of Medicine, MD, Baltimore, USA; Department of Pathology, Brigham and Women’s Hospital, Boston, MA, USA; Hacettepe University Faculty of Medicine, Department of Pathology, Ankara, Turkey; Department of Biology, Institute of Molecular Systems Biology, ETH, Zürich, Switzerland; Faculty of Science, University of Zürich, Zürich. Switzerland; Department of Urology, James Buchanan Brady Urological Institute, Johns Hopkins University School of Medicine, Baltimore, MD, USA; Institute of Basic Medical Sciences, Westlake Institute for Advanced Study, Hangzhou, Zhejiang Province, China

**Keywords:** Cell line models, castration resistance, androgen signaling, xenograft

## Abstract

**BACKGROUND:** Resistance to androgen deprivation therapies is a major driver of mortality in advanced prostate cancer. Therefore, there is a need to develop new pre-clinical models that allow the investigation of resistance mechanisms and the assessment of drugs for the treatment of castration resistant prostate cancer.

**METHODS:** We generated two novel cell line models (LAPC4-CR and VCaP-CR) which were derived by passaging LAPC4 and VCaP cells *in vivo* and *in vitro* under castrate conditions. We performed detailed transcriptomic (RNA-seq) and proteomic analyses (SWATH-MS) to delineate expression differences between castration-sensitive and castration-resistant cell lines. Furthermore, we characterized the *in vivo* and *in vitro* growth characteristics of the novel cell line models.

**RESULTS:** The two cell line derivatives LAPC4-CR and VCaP-CR showed castration resistant growth *in vitro* and *in vivo* which was only minimally inhibited by AR antagonists, enzalutamide and bicalutamide. High-dose androgen treatment resulted in significant growth arrest of VCaP-CR but not in LAPC4-CR cells. Both cell lines maintained AR expression, but exhibited distinct expression changes on the mRNA and protein level. Integrated analyses including data from LNCaP and the previously described castration resistant LNCaP-abl cells revealed an expression signature of castration resistance.

**CONCLUSIONS:** The two novel cell line models LAPC4-CR and VCaP-CR and their comprehensive characterization on the RNA and protein level represent important resources to study the molecular mechanisms of castration resistance.

## INTRODUCTION

Prostate cancer is the most common malignancy in men in the United States ^1^. Although mostly detected at a potentially curable stage, many patients experience disease progression and emergence of distant metastases. Metastatic prostate cancer accounts for around 30,000 deaths each year and therefore represents a major societal and healthcare burden ^1,2^. Androgen receptor (AR) signaling is a key oncogenic driver in prostate cancer progression and the current standard of care for treating metastatic prostate cancer involves pharmacological suppression of the AR signaling axis ^3^.

Despite initial and often profound responses to AR signaling inhibition, most patients show progression to castration-resistant prostate cancer (CRPC). Importantly, AR signaling remains active in the vast majority of CRPCs ^4,5^. Numerous resistance mechanisms to AR targeted therapies have been described which involve alterations of the *AR* gene through either mutations in the ligand binding domain, *AR* locus amplification, or expression of *AR* splice variants ^6-9^. In addition, alterations in AR-cofactors, mutations in proteins that show direct interaction with the AR such as FOXA1, MLL2, and UTX as well as upregulation of anti-apoptotic proteins such as Bcl-2 and activation of MAPK, PI3K and WNT signaling pathways have all been associated with castration resistance ^6,10-16^. Lastly, AR antagonism can result in tumors that bypass a functional requirement for AR, characterized by the loss of AR expression and other prostate luminal epithelial cell markers and in some cases gain of mesenchymal or neuroendocrine (NE) transcriptional programs ^4,5,17^.

Despite the large number of resistance mechanisms which were mostly characterized by analyzing patient samples, there is only a limited number of experimental models that are representative of CRPC. In general, the spectrum of prostate cancer cell lines is limited, and the number of cell line models has not significantly increased over the past decades ^18-20^. An increasing number of patient derived xenograft (PDX) models representative of the major molecular and phenotypic subgroups of prostate cancer have been established ^21-24^. Furthermore, organoid models propagated from PDXs, as well as directly from patient tumor samples have greatly enriched the spectrum of prostate cancer models ^25,26^.

However, these model systems are not easily amenable to large scale genetic and drug screening studies and the costs of maintaining such models can be significant, limiting their widespread use. In addition, there is a paucity of models that recapitulate the transition from androgen dependence to castration resistance and only a small number of cell line models show clinically relevant features of CRPC, such as robust growth in surgically or pharmacologically castrated mice or resistance to AR antagonists ^18,27-31^. Therefore, there is a critical need for novel cell line models of CRPC.

To help fill this need, we developed two novel CRPC cell line models which were derived from the commonly used prostate cancer cell lines, LAPC4 and VCaP ^20,32,33^. Although castration resistant sublines of VCaP and LAPC4 have been used in prior studies,^34-39^. We therefore present a comprehensive characterization of LAPC4-CR and VCaP-CR models and delineate transcriptional and proteomic differences between castration resistant and parental, androgen dependent lines. These cell line models, together with the extensive molecular profiling represent a resource for the investigation of castration resistant prostate cancer. All profiling data, as well as the cell lines are made available to the research community.

## MATERIALS AND METHODS

### Cell lines

LNCaP and VCaP cells were obtained from the American Type Culture Collection (ATCC, Manassas, VA). LAPC4 cells were a kind gift of Dr. John Isaacs, Johns Hopkins University (Baltimore, MD). LNCaP-abl cells were a gift of Dr. Zoran Culig, Innsbruck Medical University (Innsbruck, Austria) ^30^. LNCaP cells were grown in RPMI 1640 (Thermo) supplemented with 10% fetal bovine serum (FBS, Thermo) on Cell+ cell culture flask (Sarstedt, Nümbrecht, Germany). LNCaP-abl cells were grown in RPMI 1640 supplemented with 10% charcoal stripped fetal bovine serum (CSS) on Cell+ cell culture flask (Sarstedt). LAPC4 cells were grown in Iscove’s modified medium (Thermo) supplemented with 10% fetal bovine serum (FBS). LAPC4-CR cells were grown in RPMI 1640 supplemented with 10% CSS with 1x B27 (Thermo Fisher, Waltham, MA) on Cell+ cell culture flask (Sarstedt). VCaP cells were grown in DMEM (ATCC) supplemented with 10% FBS. VCaP-CR cells were grown in RPMI 1640 and 10% CSS supplemented with 1x B27 (Thermo Fisher) on Cell+ cell culture flask (Sarstedt). All cells were maintained under 5% CO2 in a humidified incubator at 37°C. Cell line authenticity and mycoplasma contamination was routinely confirmed by PCR based assays and STR genotyping, respectively, in 6-10-month intervals. Phenotypic, culturing and molecular details of all cell lines used in this study are summarized in **Supplementary Table 1**.

### Transcriptomic analysis

RNAs were extracted using the RNeasy Mini kit (Qiagen). Purified RNA was then used to prepare libraries which were sequenced on an Applied Biosystems SOLiD (V3). Sequencing reads were aligned to hg18 (NCBI36) and initially evaluated using Bioscope. Differential expression analysis was performed by DESeq on gene-level counts of properly paired reads extracted from alignment files and quantified by htseq-count with respect to features of an ensembl gene annotation file (ftp://ftp.ensembl.org/pub/release-54/gtf/homo_sapiens/Homo_sapiens.NCBI36.54.gtf.gz). Pairwise comparisons of each castrate resistant cell line versus its parent were performed, with significance assessed relative to an adjusted p-value level of 0.05 based on the Benjamini-Hochberg method. For survival analysis, we assessed 5 downregulated genes enriched in CR cell lines and mCRPC using previously published microarray gene expression dataset of primary prostate cancer (n=79) with follow-up biochemical recurrence-free survival data ^40^. The summation of average gene expression intensity of *BCHE, SPON2, GDF15, ZBTB16* and *ADAMTS1* was used as an aggregate signature score. The optimal cut-off point for the aggregate signature score was determined using the maxStat R-package and Kaplan Meier survival analysis was performed using the survival R-package. All primary expression data can be accessed on the Gene Expression Omnibus (GSE178963).

### *In vitro* cell proliferation studies

Cell growth was measured using the Incucyte® live cell imaging platform (Essen Bioscience, Ann Arbor, MI, USA). Cell surface area and percent confluence were calculated using Incucyte® Base Software package, and growth curves were plotted in Prism 8 (GraphPad Software, San Diego, CA).

### Proteomic analysis

SWATH-MS was performed as described previously ^41^. In brief, cell pellets were processed using a Barocycler NEP2320-45k (PressureBioSciences, Inc, South Easton, MA) in lysis buffer containing 8 M urea, 0.1 M ammonium bicarbonate, COMPLETE protease inhibitor cocktail (Roche), and PhosSTOP phosphatase inhibitor cocktail (Roche). Proteins were first digested using Lys-C (Wako, enzyme-to-substrate ratio 1:40) in the barocycler at 33 °C for 45 cycles, each consisting of 50 sec at 20,000 p.s.i. and 10 seconds at ambient pressure, followed by trypsin (Promega) digestions at 33 °C for 90 cycles of 50 sec at 20,000 p.s.i. and 10 sec at ambient pressure. Peptides were cleaned using SEP-PAK C18 cartridges (Waters Corp., Milford, MA) and dried under vacuum. Afterwards, they were reconstituted in HPLC grade water containing 0.1% formic acid and 2% acetonitrile and analyzed on a TripleTOF 5600 mass spectrometer (SCIEX) operated in SWATH-MS mode. Data were analyzed using OpenSWATH with a cancer cell line library as described previously ^42,43^.

### Immunohistochemistry and western blotting

Immunohistochemical studies were performed as described previously ^44^. Immuno-complexes were detected using the PowerVision+™ immunohistochemistry detection system from ImmunoVision Technologies Co (Norwell, MA, USA) with 3,3′-diaminobenzidine tetrahydrochloride (DAB) as the chromogen. After immunohistochemical staining, tissue sections were counterstained with hematoxylin. Slides were then visualized using a Nikon E400 microscope (Nikon Instruments, Melville, NY). Western blot analyses were carried out as described previously ^45^. Western blot analyses were performed as described previously ^45^. All antibodies used in the study are listed in **Supplementary Table 2**.

### Xenograft studies

All the animal experiments were performed according to protocols approved by the Animal Care and Use Committee at Johns Hopkins University. Athymic male nude mice (nu/nu, 8 weeks old) were obtained from Envigo (Huntingdon, UK). 1×10^6^ cells were resuspended in 80% Matrigel, 20% PBS and injected into the mouse flank. Caliper tumor size measurements were performed once a week and tumor volumes were calculated as described previously ^46^.

## RESULTS

### LAPC4-CR, a castration resistant subline of LAPC4

LAPC4 is an androgen dependent cell line that was originally derived from a lymph node metastasis of a human prostatic adenocarcinoma ^32^. To derive a castration resistant subline of LAPC4, we subcutaneously engrafted LAPC4 cells into intact nude mice and allowed the tumors to grow to a size of 300 mm^3^ prior to surgical castration. A tumor that progressed in size after castration was dissociated and tumor cells were plated and propagated using standard cell culture techniques. After 4 passages *in vitro*, cells were injected into the flank of a castrate male mouse and allowed to expand *in vivo* before re-establishing the line (named LAPC4-CR thereafter) *in vitro* (**Figure 1A**). Grown on standard cell culture flasks, LAPC4-CR cells showed an epithelioid morphology with cells growing adherently in small clusters (**Figure 1B**). Engraftment of LAPC4-CR cells into castrate mice showed robust tumor growth *in vivo*, establishing the castration resistant phenotype of this cell line (**Figure 1C**).

**Figure 1.**
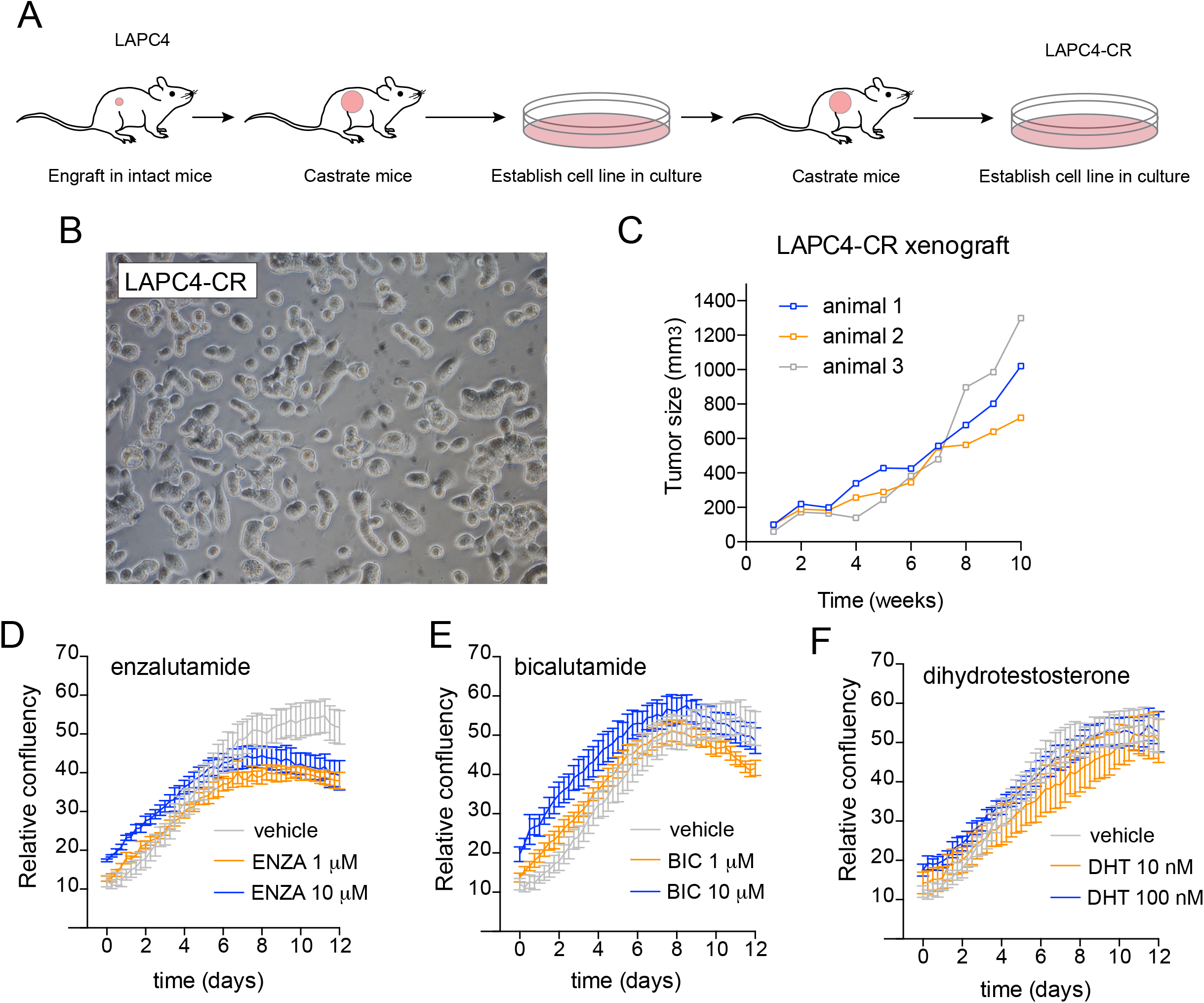
Generation of a castration resistant subline of LAPC4 (LAPC4-CR). A. Graphical summary of cell line generation. B. Representative phase contrast micrograph of LAPC4-CR cells grown *in vitro*. C. *In vivo* growth kinetics of LAPC4-CR cells engrafted in castrate nude mice. *In vitro* growth kinetic studies based on determination of cell confluency by live cell imaging of LAPC4-CR cells treated with (D) enzalutamide (vehicle control, 1μM, 10μM), (E) bicalutamide (vehicle control, 1μM, 10μM) or (F) dihydrotestosterone (DHT) (vehicle control, 10nM, 100nM).

To further characterize the growth pattern of LAPC4-CR cells in response to the AR antagonists we exposed LAPC4-CR cells to increasing doses of enzalutamide and bicalutamide or vehicle control and monitored cell growth over a 12-day period (**Figure 1D-E**). Neither treatment with enzalutamide nor bicalutamide resulted in significant changes in cell proliferation, suggesting that LAPC4-CR are intrinsically resistant to pharmacological AR pathway inhibition. Since prior reports had demonstrated a growth suppressive effect of several castration resistant prostate cancer cell lines to supraphysiological levels of androgens ^47-50^, we sought to evaluate the effect of dihydrotestosterone treatment on LAPC4-CR cell growth. Ten and 100 nM of DHT did not show any discernable effect of LAPC4-CR growth (**Figure 1F**).

### VCaP-CR, a castration resistant subline of VCaP

The VCaP cell line was derived from a vertebral prostate cancer metastasis ^33^. It shows robust *in vitro* and *in vivo* growth and is sensitive to AR pathway inhibition, despite a well-documented high-level *AR* locus amplification. To generate a castration resistant subline of VCaP, we followed a similar approach as described above for LACP4-CR. In brief, VCaP cells were engrafted into intact nude mice, which were surgically castrated at a tumor size of 300 mm^3^. Tumor tissue from a recurrent tumor was harvested and tumor cells were established to grow in 2D culture conditions resulting in the novel castration resistant VCaP subline named hereafter VCaP-CR **(Figure 2A)**. *In vitro*, VCaP-CR showed epithelioid morphology with small cells with limited cytoplasm which were adherently growing as single cells or in small clusters (**Figure 2B**). *In vivo*, the line can be propagated in castrate mice (**Figure 2C**). Similar to LAPC4-CR, VCaP-CR cells were resistant to AR antagonists (enzalutamide and bicalutamide) (**Figure 2D-EF**). However, treatment with 10 nM and 100 nM DHT resulted in profound growth suppression, suggesting that VCaP-CR are sensitive to high dose androgen treatment (**Figure 2F**).

**Figure 2.**
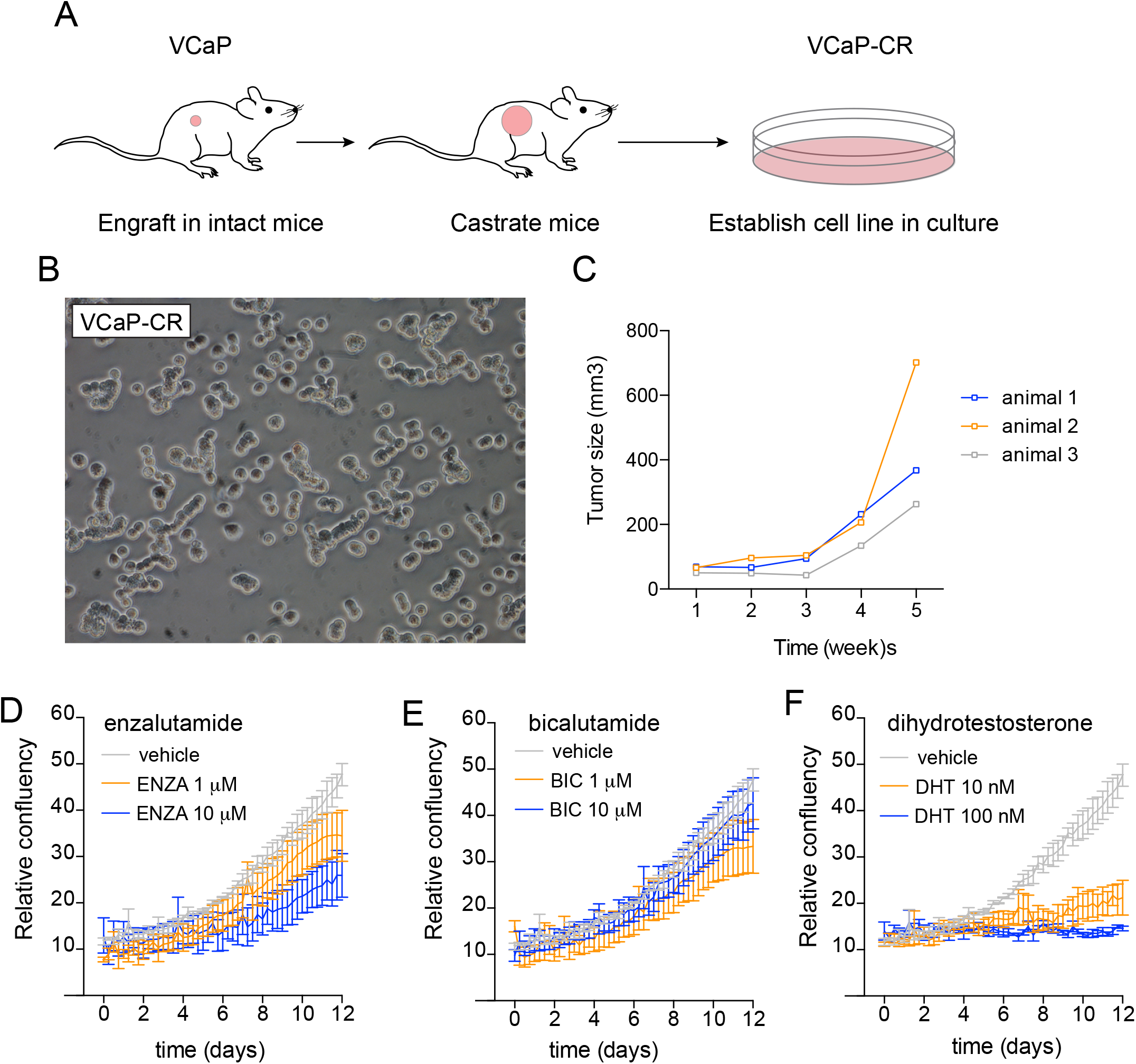
Generation of a castration resistant subline of VCaP (VCaP-CR). A. Graphical summary of cell line generation. B. Representative phase contrast micrograph of LAPC4-CR cells grown *in vitro*. C. *In vivo* growth kinetic of LAPC4-CR cells engrafted in castrate mice. *In vitro* growth kinetic studies based on determination of cell confluency by live cell imaging of LAPC4-CR cells treated with (D) enzalutamide (ENZA) (vehicle control, 1μM, 10μM), (E) bicalutamide (BIC) (vehicle control, 1μM, 10μM) or (F) DHT (vehicle control, 10nM, 100nM).

### Transcriptomic differences in cell line models of castration resistance

To determine transcriptional differences between androgen dependent and castration resistant cell lines, we performed RNA-Seq studies on the novel models described above (VCaP/VCaP-CR and LAPC4/LAPC4CR) in addition to previously described models (LNCaP/LNCaP-abl) ^20,30,51^. All expression analyses were performed with cell lines grown in their appropriate basal growth medium, which includes regular fetal bovine serum (FBS) for LNCaP, LAPC4 and VCaP and charcoal stripped FBS for LNCaP-abl, LAPC4-CR and VCaP-CR. Principal component analyses showed a relatively tight clustering of isogenic parental and castration resistant lines, but substantial differences between different cell line models (**Figure 3A**). This suggests that despite different growth phenotypes, androgen dependent and castration resistant sublines maintain similar global gene expression pattern. To investigate genes and pathways that were coordinately dysregulated in castration resistant models we performed differential expression analyses in all cell pairs (**Supplementary Table 3**). Although within a given cell line pair the number of differentially expressed genes were relatively high (155, 136 and 51 for LNCaP/LNCaP-abl, LAPC4/LAPC4CR and VCaP/VCaP-CR respectively; **Figure 3B, Table 1, Supplementary Table 2**) the intersection of these gene lists showed a greatly reduced number of shared expression changes that were common to all three models (**Figure 3B, Table 1**). Importantly, differentially expressed genes were enriched for known androgen regulated genes ^52^ (**Supplementary Table 4, see Supplementary Table 5 for full gene list**), but also comprised numerous genes without prior evidence for androgen regulation. This suggests that AR-dependent and independent signaling pathways are altered in castration resistant models. Indeed, pathway analyses revealed several gene sets involved in actin binding, cytoskeleton, alternative splicing and protein binding to be up regulated in all castration resistant cell line models (**Supplementary Table 6**). To determine if genes differentially expressed in cell line models were also transcriptionally altered in clinical mCRPC samples, we queried a publicly available dataset comprised of primary hormone naive tumors and mCRPCs (**Figure 3C**) ^10^. Although directionality and magnitude of expression changes between primary tumors and mCRPC varied for individual genes, we observed significantly lower expression in mCRPC for a subset of genes, including *BCHE, SPON2, GDF15, ZBTB16* and *ADAMTS1*, showing differential expression in castration resistant cell lines. Interestingly, Kaplan-Meier analyses revealed that primary tumors expressing low levels of these five genes showed earlier biochemical recurrence (P = 0.007) (**Figure 3D**). Collectively these data suggest that differential expression pattern observed in cell line models can also be seen in clinical mCRPC samples, therefore establishing the translational relevance of these novel cell line models.

**Figure 3.**
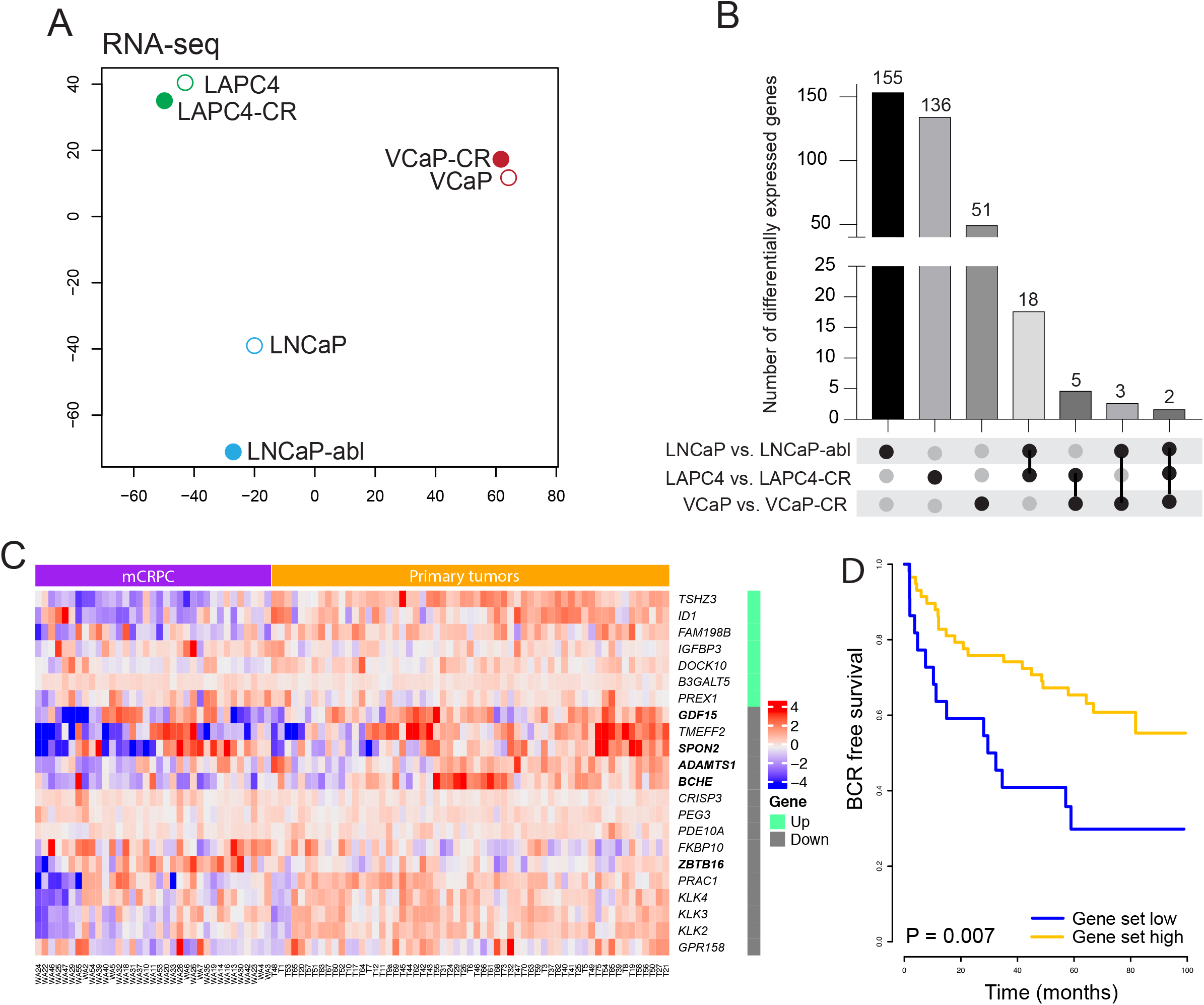
Transcriptional profiling of androgen dependent and castration resistant prostate cancer cell lines. A. Principal component analysis of RNA-seq data from LNCaP, LNCaP-abl, LAPC4, LAPC4-CR, VCaP, VCaP-CR reveals clustering of cell line pairs according to their parental origin. B. Bar graph showing number of statistically significant (P < 0.05) differentially expressed genes in comparisons between parental and castration resistant lines; line segments between the black dots indicate the intersection of differentially expressed genes for the comparisons marked by the black dots. C. Heatmap shows expression of genes in primary prostate cancers and mCRPC samples from a previously published cohort with differential expression in parental and castration resistant cells in at least 2 models (also see **Table 1**)^10^. D. Kaplan Meier plot showing time to biochemical recurrence estimates derived from primary tumor analysis of a publicly available dataset ^40^ stratified by high (orange) or low (blue) expression of *BCHE, SPON2, GDF15, ZBTB16* and *ADAMTS1* (all down regulated genes in castration resistant cell lines). Note, genes used in the Kaplan Meier model are printed in bold in panel C.

**Table 1.**
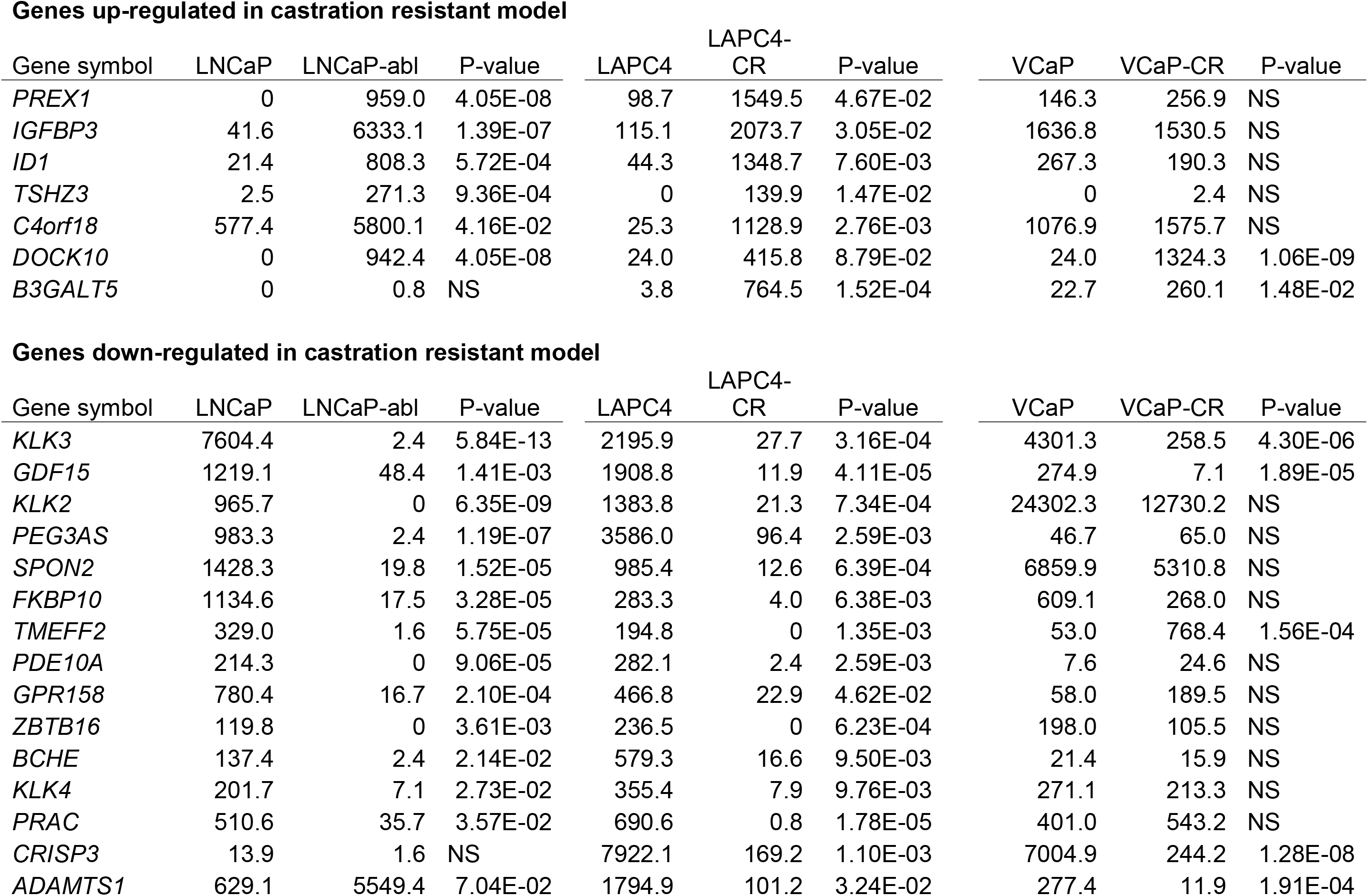
Differentially expressed genes in CRPC cell lines. Genes with up- or down-regualtion in CRPC models. Numbers indicate normalized Reads Per Kilobase of transcript per Million reads mapped (RPKM).

| Gene symbol | LNCaP | LNCaP-abl | P-value | LAPC4 | LAPC4-CR | P-value | VCaP | VCaP-CR | P-value |
| --- | --- | --- | --- | --- | --- | --- | --- | --- | --- |
| *PREX1* | 0 | 959.0 | 4.05E-08 | 98.7 | 1549.5 | 4.67E-02 | 146.3 | 256.9 | NS |
| *IGFBP3* | 41.6 | 6333.1 | 1.39E-07 | 115.1 | 2073.7 | 3.05E-02 | 1636.8 | 1530.5 | NS |
| *ID1* | 21.4 | 808.3 | 5.72E-04 | 44.3 | 1348.7 | 7.60E-03 | 267.3 | 190.3 | NS |
| *TSHZ3* | 2.5 | 271.3 | 9.36E-04 | 0 | 139.9 | 1.47E-02 | 0 | 2.4 | NS |
| *C4orf18* | 577.4 | 5800.1 | 4.16E-02 | 25.3 | 1128.9 | 2.76E-03 | 1076.9 | 1575.7 | NS |
| *DOCK10* | 0 | 942.4 | 4.05E-08 | 24.0 | 415.8 | 8.79E-02 | 24.0 | 1324.3 | 1.06E-09 |
| *B3GALT5* | 0 | 0.8 | NS | 3.8 | 764.5 | 1.52E-04 | 22.7 | 260.1 | 1.48E-02 |

**Genes down-regulated in castration resistant model**
| Gene symbol | LNCaP | LNCaP-abl | P-value | LAPC4 | LAPC4-CR | P-value | VCaP | VCaP-CR | P-value |
| --- | --- | --- | --- | --- | --- | --- | --- | --- | --- |
| *KLK3* | 7604.4 | 2.4 | 5.84E-13 | 2195.9 | 27.7 | 3.16E-04 | 4301.3 | 258.5 | 4.30E-06 |
| *GDF15* | 1219.1 | 48.4 | 1.41E-03 | 1908.8 | 11.9 | 4.11E-05 | 274.9 | 7.1 | 1.89E-05 |
| *KLK2* | 965.7 | 0 | 6.35E-09 | 1383.8 | 21.3 | 7.34E-04 | 24302.3 | 12730.2 | NS |
| *PEG3AS* | 983.3 | 2.4 | 1.19E-07 | 3586.0 | 96.4 | 2.59E-03 | 46.7 | 65.0 | NS |
| *SPON2* | 1428.3 | 19.8 | 1.52E-05 | 985.4 | 12.6 | 6.39E-04 | 6859.9 | 5310.8 | NS |
| *FKBP10* | 1134.6 | 17.5 | 3.28E-05 | 283.3 | 4.0 | 6.38E-03 | 609.1 | 268.0 | NS |
| *TMEFF2* | 329.0 | 1.6 | 5.75E-05 | 194.8 | 0 | 1.35E-03 | 53.0 | 768.4 | 1.56E-04 |
| *PDE10A* | 214.3 | 0 | 9.06E-05 | 282.1 | 2.4 | 2.59E-03 | 7.6 | 24.6 | NS |
| *GPR158* | 780.4 | 16.7 | 2.10E-04 | 466.8 | 22.9 | 4.62E-02 | 58.0 | 189.5 | NS |
| *ZBTB16* | 119.8 | 0 | 3.61E-03 | 236.5 | 0 | 6.23E-04 | 198.0 | 105.5 | NS |
| *BCHE* | 137.4 | 2.4 | 2.14E-02 | 579.3 | 16.6 | 9.50E-03 | 21.4 | 15.9 | NS |
| *KLK4* | 201.7 | 7.1 | 2.73E-02 | 355.4 | 7.9 | 9.76E-03 | 271.1 | 213.3 | NS |
| *PRAC* | 510.6 | 35.7 | 3.57E-02 | 690.6 | 0.8 | 1.78E-05 | 401.0 | 543.2 | NS |
| *CRISP3* | 13.9 | 1.6 | NS | 7922.1 | 169.2 | 1.10E-03 | 7004.9 | 244.2 | 1.28E-08 |
| *ADAMTS1* | 629.1 | 5549.4 | 7.04E-02 | 1794.9 | 101.2 | 3.24E-02 | 277.4 | 11.9 | 1.91E-04 |

### Immunohistochemical assessment of key signaling pathways

To investigate the expression of key proteins relevant for prostate cancer biology, we performed immunohistochemical expression analyses on LAPC4/LAPC4-CR and VCaP/VCaP-CR cells (**Figure 4**). We observed that, although all cell lines expressed AR, LAPC4-CR cells showed significantly reduced AR levels, which was accompanied by a reduced expression of the AR target NKX3.1 (**Figure 4**). Neuroendocrine markers such as synaptophysin (SYN) and chromogranin B (CHGA) were expressed in VCaP and VCaP-CR cells, but not in LAPC4 cells. This supported the prior observation that VCaP cells should be classified as an amphicrine cell line, characterized by co-expression of AR and neuroendocrine markers ^5^. In addition, VCaP and VCaP-CR cells also showed expression of the glucocorticoid receptor (GR) in a subset of cells (GR). GR expression was previously shown to be increased in castration resistant cell line models and mCRPC tissues ^53^. Interestingly however, GR expression appeared to be higher in parental VCaP cells compared to VCaP-CR cells. Furthermore, VCaP cells, which are known to harbor a *TMPRSS2-ERG* rearrangement showed high levels or ERG expression which was maintained in VCaP-CR cells ^54^. The pioneer transcription factor FOXA1 showed uniformly high expression in VCaP and VCaP-CR cells but was present at relatively lower levels in LAPC4 and LAPC4-CR. Both LAPC4 and VCaP cell lines showed retained PTEN and RB1 expression and high levels of nuclear cMYC staining (**Supplementary Figure 1**). In addition, all cell line models showed nuclear p53 accumulation consistent with the known *TP53* mutations in LAPC4 (R175H) and VCaP (R248W) (**Supplementary Figure 1**) ^20^.

**Figure 4.**
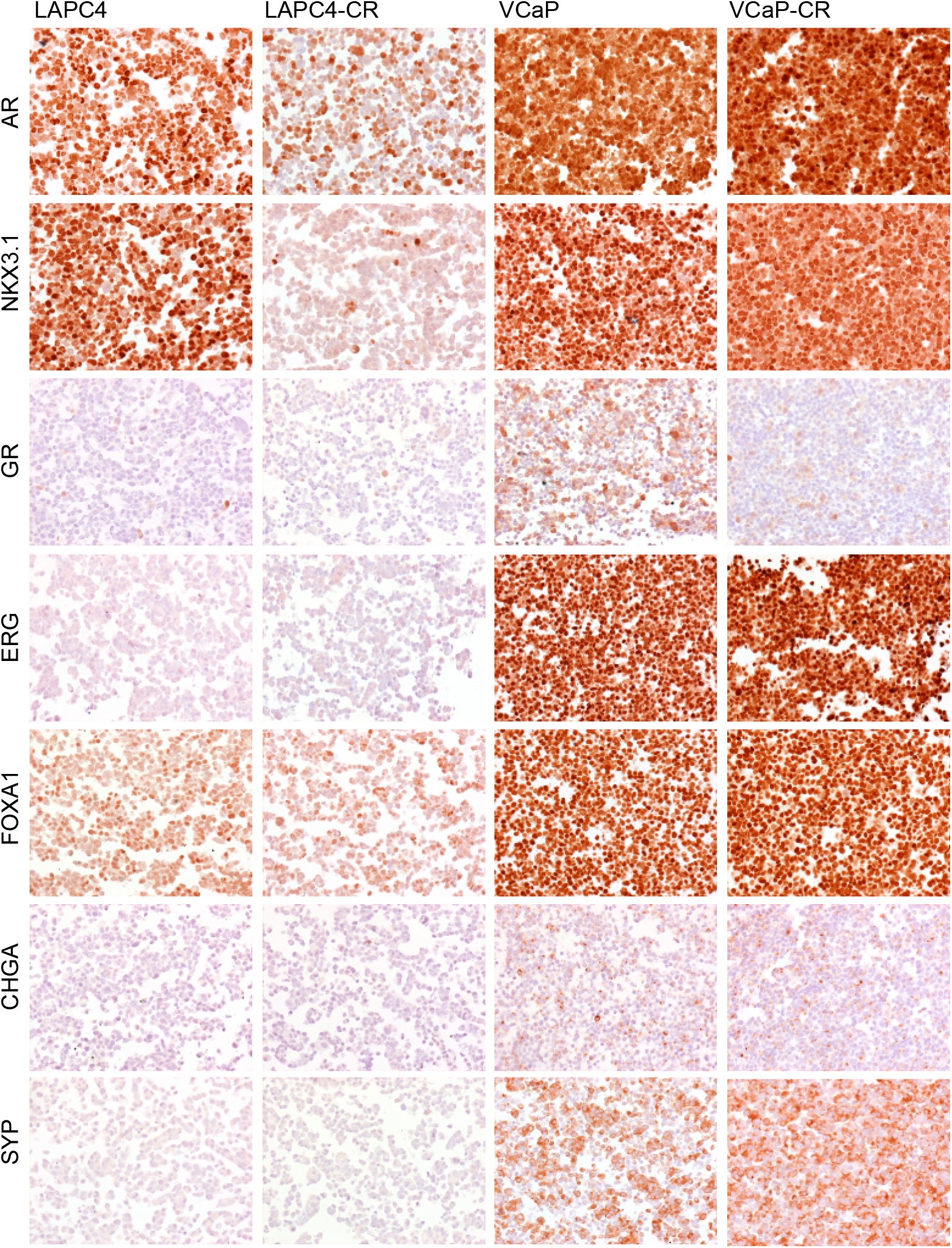
Immunohistochemical phenotyping of LAPC4, LAPC4-CR, VCaP and VCaP-CR cells. Representative micrographs showing immunohistochemical staining for AR, NKX3.1, GR, ERG, FOXA1, chromogranin A (CHGA) and synaptophysin (SYP) in formalin fixed paraffin embedded LAPC4, LAPC4-CR, VCaP and VCaP-CR cells.

### Proteomic differences in cell line models of castration resistance

To probe the expression of key signaling pathways, we performed immunoblotting experiments on LNCaP, LNCaP-abl, LAPC4, LAPC4-CR, VCaP and VCaP-CR cells (**Figure 5A**). AR levels appeared to be modestly decreased in castration resistant models. In addition to full length AR, VCaP and VCaP-CR cells also showed expression of a lower molecular weight AR variant, consistent with the previously reported AR-V7 ^9^. Differences in AR expression levels were accompanied by decreased expression of the AR target gene NKX3.1 in LNCaP-abl and LAPC4-CR suggesting reduced AR signaling activity. VCaP-CR cells however showed high NKX3.1 expression indicating a ligand independent activity of AR in these cells. To probe for the activity of AKT and MAPK signaling, we performed immunoblotting for phospho-p44/42 MAPK and phospho-AKT. Interestingly, we observed a strong increase in MAPK phosphorylation in LAPC4-CR and VCaP-CR cells (**Figure 5A**). Whereas both LNCaP and LNCaP-abl cells showed high levels of AKT phosphorylation, LAPC4-CR cells showed increased AKT phosphorylation compared to parental LAPC4 cells suggesting a potential activation of AKT and MAPK signaling during the transition to castration resistance.

**Figure 5.**
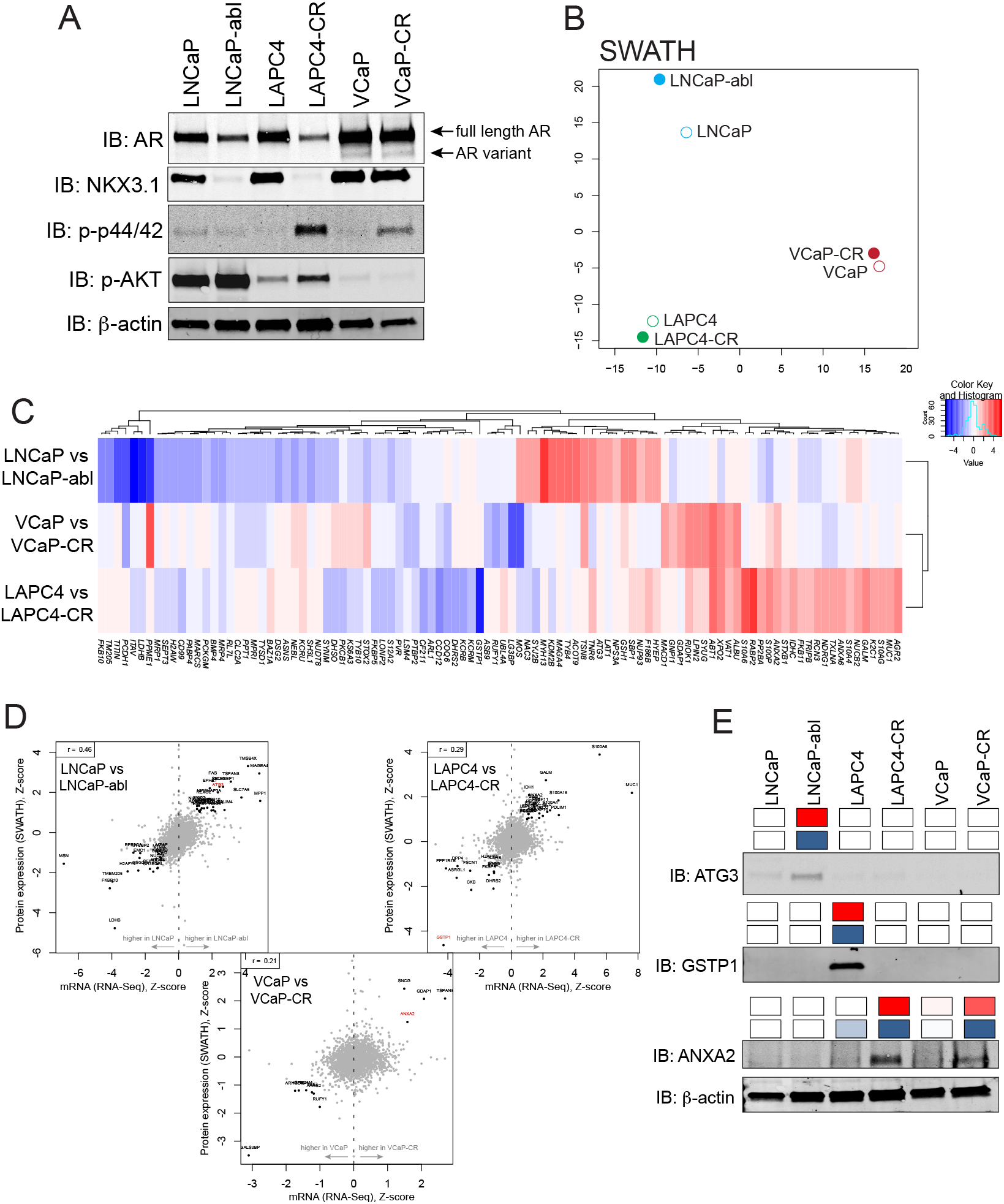
Proteomic profiling of androgen dependent and castration resistant prostate cancer cell lines. A. Western blot analysis showing AR and NKX3.1 expression as well as abundance of phospho-p44/42 MAPK (Erk1/2) (Thr202/Tyr204) and Phospho-Akt (Ser473) in LNCaP, LNCaP-abl, LAPC4, LAPC4-CR, VCaP, VCaP-CR. B. Principal component analysis of SWATH-MS data from LNCaP, LNCaP-abl, LAPC4, LAPC4-CR, VCaP, VCaP-CR reveals clustering of cell line pairs according to their parental origin. C. Heatmap of paired comparisons of differentially expressed proteins in parental and castration resistant cell line, red indicates higher expression in castration resistant model, blue indicates higher expression in the parental line. D. Correlation between mRNA (x-axis) and protein (y-axis) expression in parental and castration resistant cell line pairs. Names of significantly differentially expressed proteins are indicated. Proteins printed in red are further validated by western blot analysis. E. Differential expression of ATG3, GSTP1 and ANXA2 in LAPC4, LAPC4-CR, VCaP and VCaP-CR cells. Bands show immunoreactivity in western blot. Boxes (top row) indicate mRNA expression based on RNA-seq (red = high expression, white low/no expression); boxes (second row) indicate protein expression based on SWATH-MS (blue = high expression, white low/no expression). Beta-actin serves as loading control.

To further characterize global proteomic differences between androgen dependent and castration resistant cell line models we performed sequential window acquisition of all theoretical fragment-ion mass spectrometry (SWATH-MS) on LNCaP, LNCaP-abl, LAPC4, LAPC4-CR, VCaP and VCaP-CR cells as described previously ^41^. Using this approach, we identified >3100 proteins in each sample. Principal component analyses revealed clustering of cell lines based on their parental origin (**Figure 5B, Supplementary Table 7**). This finding is reminiscent of the principal component analyses based on RNA-seq data (**Figure 3A**) and suggest that despite the phenotypic conversion to castration resistance, there is limited global change in protein expression between the different cell line models. Proteins uniformly downregulated in all CR cell lines included MRP4 (encoded by *ABCC4*) and PVR. Although there were no proteins upregulated in all CR models, ALBU, VAT1, XPO2 and ABT1 were overexpressed in VCaP-CR and LAPC4-CR relative to their corresponding parental lines (**Figure 5C, Supplementary Table 7**). The datasets generated as part of this study allows the correlation of different data streams, which might provide new insights into the transcriptional and translational regulation as well as protein homeostasis. To showcase the potential application of the dataset, we correlated mRNA expression determined by RNA-seq with protein abundance determined by SWATH-MS. Although mRNA and protein levels were correlated, the correlation coefficients were generally low (0.21 to 0.46, **Figure 5D**) and varied between different cell line models. These findings are in line with prior reports showing that quantitative assessment of protein and mRNA abundance can show differences and highlight the complex regulation of biological pathways at different levels ^43,55,56^. To corroborate proteins which showed differential expression, we performed western blot analysis for ATG3 (Autophagy related 3; differentially expressed in LNCaP and LNCaP-abl), GSTP1 (Glutathione S-transferase pi; differentially expressed in LAPC4 and LAPC4-CR) and ANXA2 (Annexin A2; differentially expressed in LAPC4 and LAPC4-CR and VCaP and VCaP-CR) (**Figure 5E**). Immunoreactivity pattern on western blots confirmed the differential expression observed by SWATH-MS and RNA-seq. More broadly, these data show the robustness of the measurements presented here and validate the findings of our unbiased protein and mRNA expression analyses.

## DISCUSSION

Prostate cancer has been recognized as an androgen sensitive and androgen driven disease ^3,57^. Therapeutic strategies that interfere with the gonadal/extragonadal production of androgens or the action of the androgen receptor itself have been established as a standard of care for patients with metastatic disease. Despite initial and often profound responses to this hormonal therapy, most patients show progression to CRPC ^2,3,58^. Castration resistance remains the major challenge in the management of patients with advanced prostate cancer. Although several resistance mechanisms have been recognized, the biology of castration resistant disease is complex. The lack of a representative number of model systems that enable the investigation of processes involved in the conversion from androgen dependence to castration resistance has also hindered progress of discovery in this space.

Amongst the limited number of AR responsive prostate cancer cell lines, LNCaP, VCaP, and LAPC4 are the most commonly used ^18-20^. Although all three lines were derived from patients with metastatic prostate cancer with a prior treatment history that included androgen deprivation therapies, all 3 lines continue to show an androgen dependent growth phenotype. Prior studies have developed derivatives of these cell lines that grow in castrate mice or under androgen deprived media conditions *in vitro* ^18,19,32,59^. The majority of these CRPC sublines were derived from LNCaP cells ^18,51^,including C4-2, C4-2B, LNCaP-AI, LNCaP-abl and LNCaP95 ^27-31^. These lines show variable *in vivo* and *in vitro* growth characteristics and molecular changes ^52,60^. Here, we describe two novel cell lines which represent castration resistant sublines of the commonly used, androgen dependent prostate cancer cells lines LAPC4 and VCaP. It is worth noting that other groups have described similar models previously ^34-39,61^. However, for many of these lines there is limited profiling data publicly available. We therefore aimed to generate new cell line models and associated profiling data as a resource for the prostate cancer research community. We hope that these models will represent valuable tools for the discovery of molecular alterations associated with castration resistance.

LAPC4-CR and VCaP-CR cells both grow under castration conditions *in vitro* and *in vivo* and show resistance to pharmacological inhibition of the AR. Interestingly however, their response to dihydrotestosterone differs. Whereas LAPC4-CR cells show no discernable difference in in vitro growth in response to DHT, VCaP-CR cells exhibits a profound growth suppression at 10 and 100 nM of DHT. This paradoxical growth suppressive effect to supraphysiological concentrations of androgens has been described for other cell line models and has led to the clinical evaluation of high dose testosterone therapies in men with castration resistant prostate cancer ^49^. Several clinical trials have shown promising results with profound PSA and radiographic responses in a subset of patients receiving testosterone therapy ^62-65^. Although the mechanism underlying growth suppressive effect of testosterone is unclear, a recent study has suggested that high AR expression levels and amplification of the AR locus are associated with increased androgen-induced growth suppression ^47^. VCaP and VCaP-CR show the highest level for AR protein expression and are known to harbor AR amplification which could contribute to the profound effects in growth suppression upon androgen treatment seen in these models.

Although the conversion to castration resistance was associated with transcriptional changes and alterations in protein expression, the broad transcriptional output of androgen dependent cell lines did not differ significantly from the parental lines. In principal component analyses castration resistant cell lines clustered with their parental lines suggesting a closer similarity of expression changes between cell lines originating from the same parental clone than cell lines with a castration resistant phenotype. This suggests that expression changes associated with castration resistance can vary greatly between different cell line models. This also reflects the results from in depth analysis of clinical mCRPC specimens, which show a high level of inter-individual heterogeneity in expression profiles ^5^ and suggests that the conversion to castration resistance likely involves distinct subsets of genes and not a global transcriptional reprogramming. In this context it is worth noting that most expression changes observed between parental and CR lines were private and therefore only found in one of the cell line pairs. KLK3 and GDF15 were the only two genes that were downregulated in all 3 castration resistant models relative to the parental line. *KLK3* encodes for prostate specific antigen (PSA) and is a known AR regulated gene. Given the fact that all castration resistant lines are cultured in the absence of androgens, it is not surprising to see KLK3 downregulation. GDF15 is a stress induced cytokine and part of the transforming growth factor beta superfamily. Its expression has been associated with cancer progression and bone metastasis formation, but the role of GDF15 in prostate cancer is poorly understood ^66,67^. GDF15 does not appear to be androgen regulated and its transcriptional control is not well studied. Although DOCK10 does not meet the statistical threshold for differential expression in all 3 models, there is a strong trend toward higher expression in the castration resistant models. DOCK10 belongs to the DOCK (Dedicator of cytokinesis protein) gene family and appears to be involved in the regulation of small G proteins (such as Cdc42) which are important for the regulation of cell migration and invasion ^68^. Other genes that are increased in expression in castration resistant models, including *PREX1, IGFBP3* and *ID1*, have recently been implicated in prostate cancer biology ^69-71^. Of particular interest, high expression of *ID1*, as seen in LNCaP-abl and LAPC4-CR cells has been shown to be associated with castration resistance through activation of epidermal growth factor receptor signaling ^72^. More broadly, these data provide evidence that alterations observed in the cell line models presented here can inform future mechanistic and translational studies.

## CONCLUSIONS

Here we describe novel cell line models of castration resistant prostate cancer and characterize their phenotype in *in vitro* and *in vivo* experiments. We demonstrate that these cell lines at least partially recapitulate changes observed in clinical samples of mCRPC suggesting that these models can be used to potentially unmask novel biological features of advanced prostate cancer. The comprehensive transcriptomic and proteomic characterization of the cell lines presented here should serve as a useful resource to study intrinsic pathways for castration resistant prostate cancer and to develop new therapies for CRPC.

## Grant Acknowledgements

This work was supported by the NIH/NCI (P50CA097186, P50CA58236, U01 CA196390, P30 CA006973, R01CA183965), the U.S. Department of Defense Prostate Cancer Research Program (W81XWH-20-1-0111, W81XWH-18-1-0406, W81XWH-18-2-0015), the Swiss National Science Foundation (#31003A_166435), the Prostate Cancer Foundation, the Safeway Foundation, the Commonwealth Foundation and the Irving Hansen Memorial Foundation.

The authors would like to thank the Sidney Kimmel Comprehensive Cancer Center Cell Imaging Facility, in particular Lillian Dasko-Vincent. We also thank Drs. John T. Isaacs and Alan K Meeker for helpful comments and discussions.

## SUPPLEMENTARY FIGURES

**Supplementary Figure 1. Additional immunohistochemical phenotyping of LAPC4, LAPC4-CR, VCaP and VCaP-CR**. LAPC4, LAPC4-CR, VCaP and VCaP-CR cells show uniform expression of PTEN, MYC and RB. All cell lines exhibit nuclear accumulation of p53 suggestive of TP53 missense mutations.

## SUPPLEMENTARY TABLES

Supplementary Table 1. Summary of characteristics of cell lines used in this study. Supplementary Table 2. Antibodies used in the study.

Supplementary Table 3. Differentially expressed genes in cell line models.

Supplementary Table 4. AR regulation of genes with significant differential expression in at least 2 cell line pairs.

Supplementary Table 5. RNA-seq gene expression results.

Supplementary Table 6. Gene set enrichment analyses with genes differentially expressed in castration resistant cell line models.

Supplementary Table 7. SWATH-MS protein expression results.

