## Supplementary Figure 1 for "PHENOTYPIC CHARACTERIZATION OF TWO NOVEL CELL LINE MODELS OF CASTRATION RESISTANT PROSTATE CANCER"

Haffner et al. Figure 1.

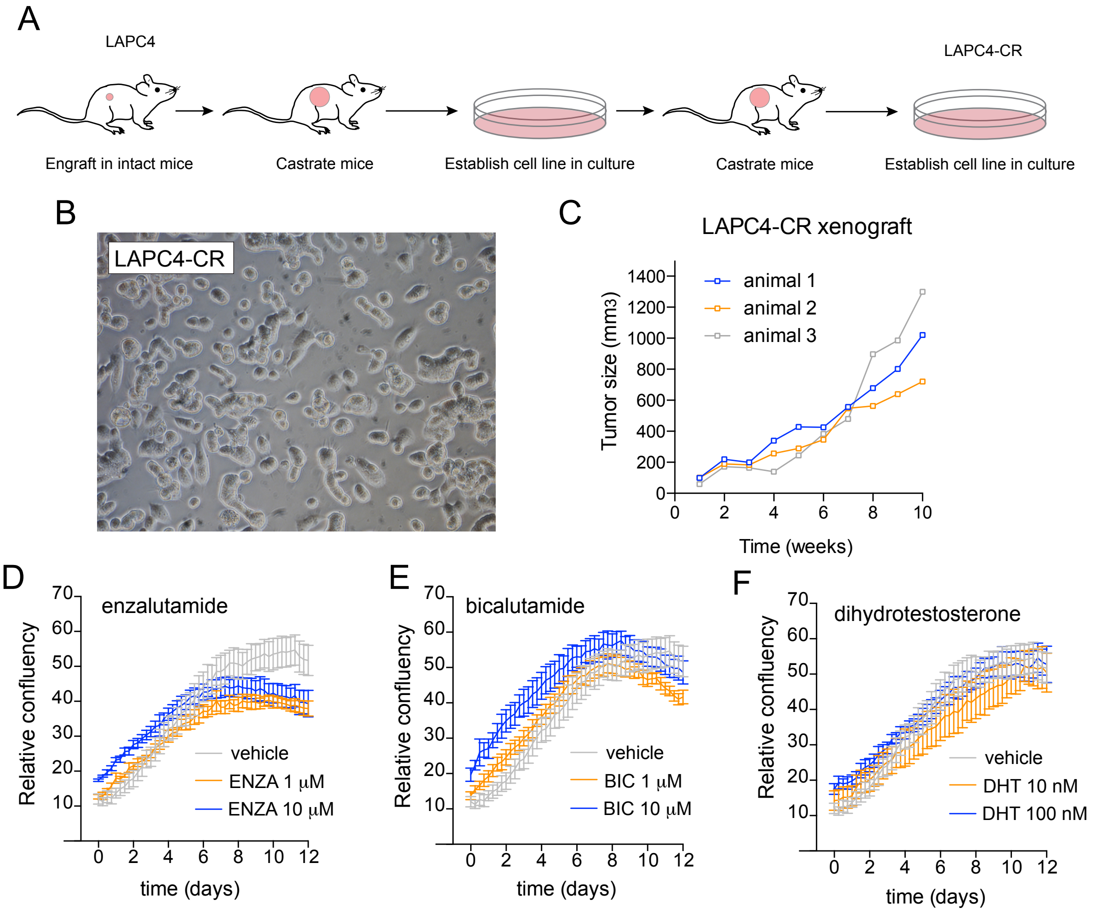

Haffner et al. Figure 2.

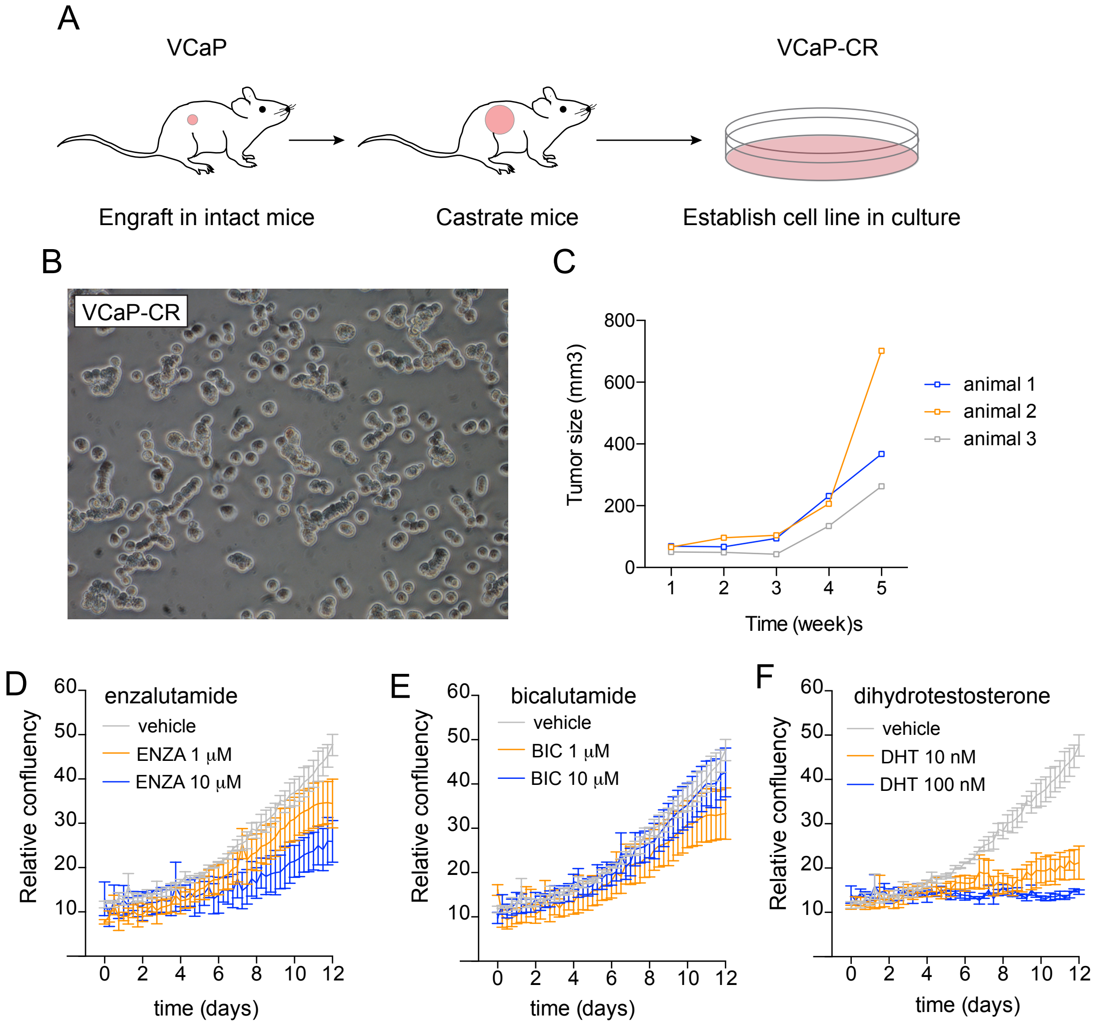

Haffner et al. Figure 3.

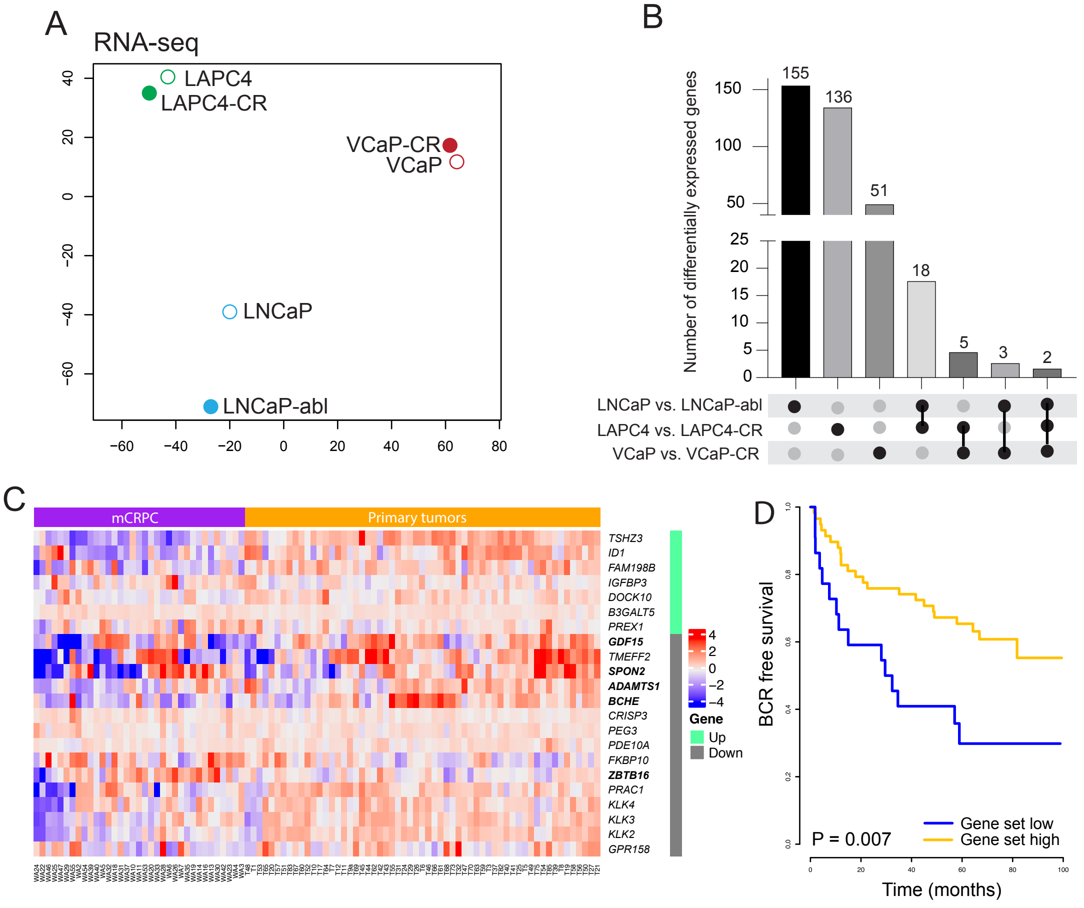

Haffner et al. Figure 4.

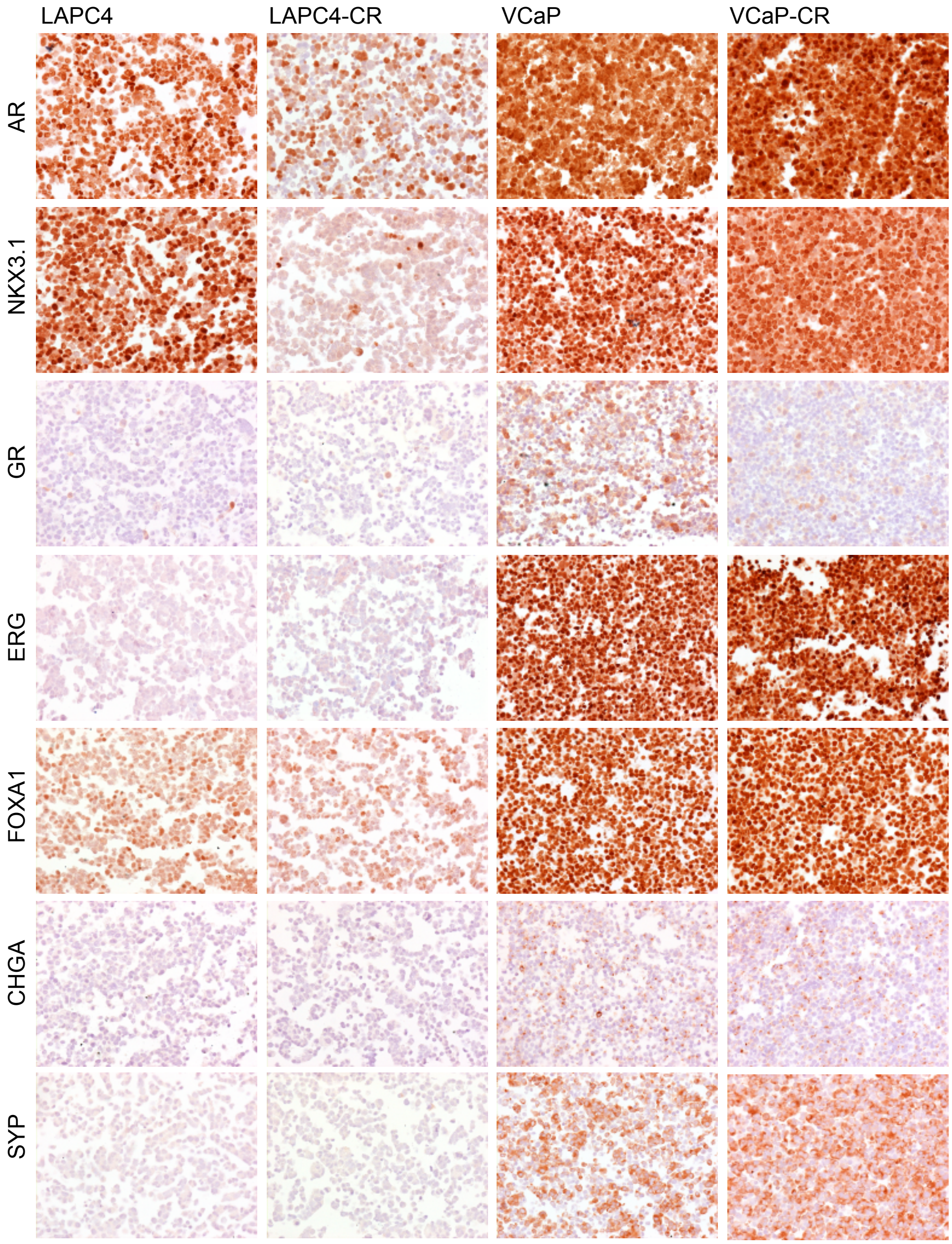

Haffner et al. Figure 5.

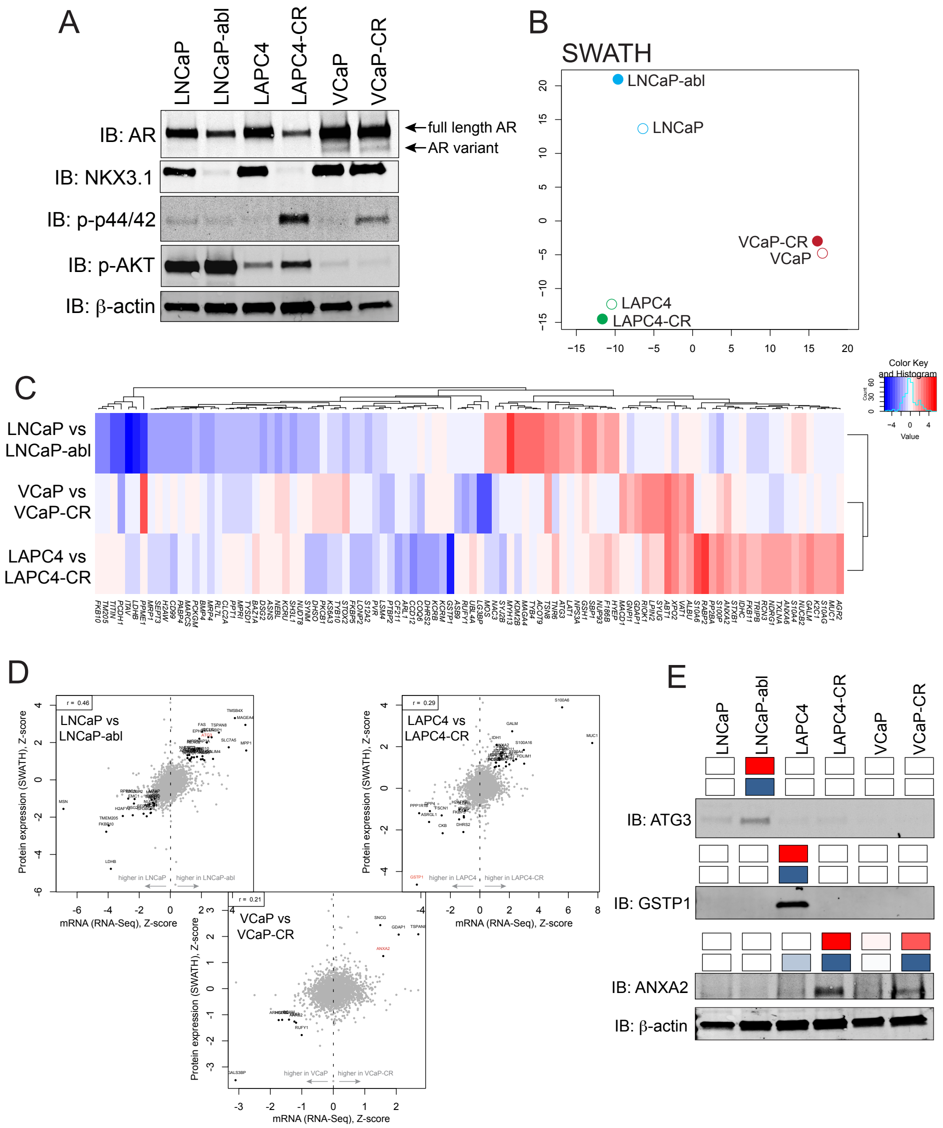

Haffner et al. Supplementary Figure 1.

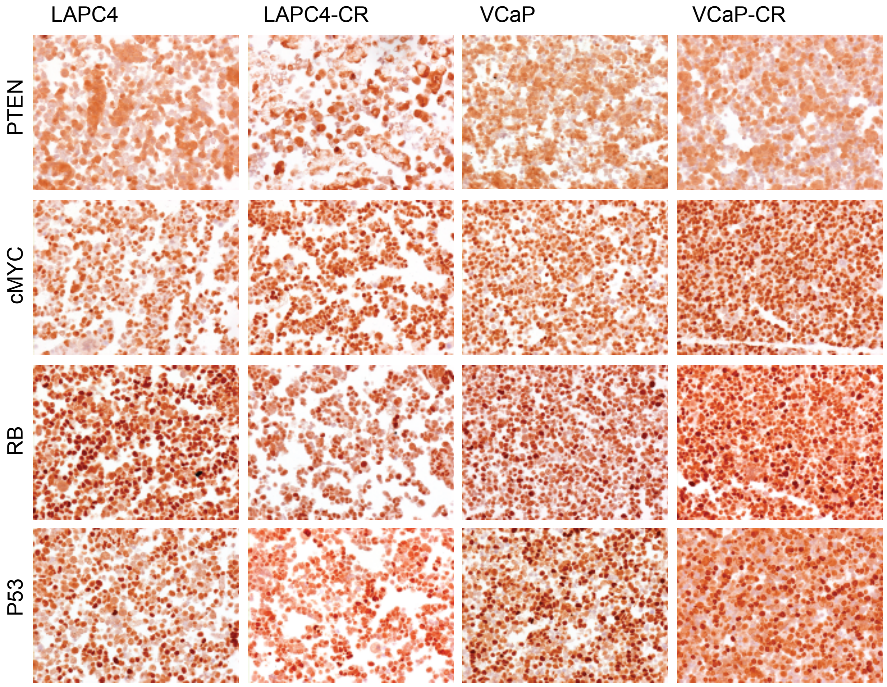
